# Bird size with dinosaur-level cancer defences: can evolutionary lags during miniaturisation explain cancer robustness in birds?

**DOI:** 10.1101/2020.10.22.345439

**Authors:** E. Yagmur Erten, Marc Tollis, Hanna Kokko

**Affiliations:** Department of Evolutionary Biology and Environmental Studies, University of Zurich, Winterthurerstrasse 190, CH-8057 Zurich, Switzerland; Arizona Cancer Evolution Center, Arizona State University, Tempe, Arizona, United States; School of Informatics, Computing, and Cyber Systems, Northern Arizona University, Flagstaff, Arizona, United States

**Keywords:** life history, Peto’s paradox, miniaturisation, maladaptation, macroevolution

## Abstract

An increased appreciation of the ubiquity of cancer risk across the tree of life means we also need to understand the more robust cancer defences some species seem to have. Peto’s paradox, the finding that large-bodied species do not suffer from more cancer even if their lives require far more cell divisions than those of small species, can be explained if large size selects for better cancer defences. Since birds live longer than non-flying mammals of an equivalent body size, and birds are descendants of moderate-sized dinosaurs, we ask whether ancestral cancer defence innovations are retained if body size shrinks in an evolutionary lineage. Our model derives selection coefficients and fixation events for gains and losses of cancer defence innovations over macroevolutionary time, based on known relationships between body size, intrinsic cancer risk, extrinsic mortality (modulated by flight ability) and effective population size. We show that evolutionary lags can, under certain assumptions, explain why birds, descendants of relatively large bodied dinosaurs, retain low cancer risk. Counterintuitively, it is possible for a bird to be ‘too robust’ for its own good: excessive cancer suppression can take away from reproductive success. On the other hand, an evolutionary history of good cancer defences may also enable birds to reap the lifespan-increasing benefits of other innovations such as flight.

## Introduction

Cancer is likely ubiquitous in nature, having been observed in all but few lineages of multicellular organisms (Aktipis et al., 2015; Albuquerque et al., 2018). Multicellularity relies on cooperation between the cells of an organism, and mutations that disrupt this cooperation can cause uncontrolled cell proliferation and oncogenesis (Domazet-Lošo and Tautz, 2010; Aktipis et al., 2015). Each cell division carries a risk of mutations. Since larger and longer-lived organisms need more divisions to reach and maintain their adult body size, the ‘all else being equal’ expectation is a higher cancer risk if either body size or lifespan increases. As such, cancer can become a limiting factor to a further evolutionary increase of either (Kokko and Hochberg, 2015). Studies in humans (Albanes et al., 1988; Green et al., 2011; Nunney, 2018) and dogs (Fleming et al., 2011; Nunney, 2013) suggest that larger individuals within a species have an increased cancer incidence. However, the between-species pattern is different: large body size and long lifespan do not combine to predict elevated cancer incidence (Abegglen et al., 2015). This lack of support for a theoretically clear a priori expectation is known as ‘Peto’s Paradox’ (Peto, 1977; Nunney, 1999).

Current comparative oncology efforts to understand cancer across the tree of life are based on the insight that species differ in past selection to improve cancer suppression mechanisms. Such selection may be stronger in large-bodied and/or long-lived species (Leroi et al., 2003; Caulin and Maley, 2011), and these species make fitting targets for understanding the molecular bases of cancer defences (Tollis et al., 2017; Sulak et al., 2016; Seluanov et al., 2018; Attaallah et al., 2020). For instance, higher rates of p53-mediated apoptosis in response to DNA damage in elephant cells are associated with multiple copies of the tumor suppressor gene TP53 in the elephant genome (Abegglen et al., 2015). Evidence also comes from smaller but unusually long-lived organisms: the long and relatively cancer-free life of naked mole rats (*Heterocephalus glaber*) has been attributed to their skin cells showing a high degree of contact inhibition which provides cancer resistance — important in a species in which reproductives, so-called queens, have exceptional longevity for their body size (Tian et al., 2013).

Lifespan correlates strongly with body size, with larger animals typically living longer (Blueweiss et al., 1978; Lindstedt and Calder III, 1981; Ricklefs, 2010). Yet, there are tantalizing differences between taxa: for their (usually small) body size, birds are relatively long lived (Partridge and Barton, 1993; Holmes and Austad, 1994; Holmes and Ottinger, 2003; Healy et al., 2014), and compared with mammals, they enjoy a delayed onset of senescence (Jones et al., 2008; Péron et al., 2010). Ages beyond 60 years old can be reached by some species. Flight has been argued to be a key innovation (Partridge and Barton, 1993) that allows birds to experience lowered extrinsic mortality from predation, a pattern repeated in flying mammals (Holmes and Austad, 1994; Healy et al., 2014; Wilkinson and Adams, 2019) which, too, live long for their body size. Obviously, a ‘slow’ life history in a flying organism can only reap the benefit of a potentially long life if its body is robust enough to escape senescence (for some time) too. One subtask within senescence avoidance is to avoid succumbing to age-related diseases such as cancer. Theory and empirical data suggest cancer incidence increases with age (Armitage and Doll, 1954; Calabrese and Shibata, 2010; Martincorena and Campbell, 2015), making the relatively low cancer incidence of birds compared with mammals and reptiles (Effron et al., 1977; Møller et al., 2017) particularly intriguing.

All extant birds belong to a lineage that shrunk from large theropod ancestors through a process of continuous miniaturisation, accompanied by innovations that eventually led to flight (Benson et al., 2014; Lee et al., 2014). Comparative studies show that bird cells are more stress resistant than rodent cells (Ogburn et al., 1998; Harper et al., 2011), but the interpretation is challenging: as the elephant example shows, more (rather than less) cell-level death might improve longevity at the organismal level. As a whole, avian cancer suppression mechanisms remain unexplored. They are particularly intriguing because an a priori beneficial trait — here, any form of cancer defence — may lose importance when body size shrinks (Kokko and Hochberg, 2015). Based on the prevalence of tumours across the tree of life (Aktipis et al., 2015), it is parsimonious to assume cancer was also a problem affecting dinosaurs (for direct evidence see Rothschild et al., 1999, 2003, 2020; Ekhtiari et al., 2020). The ancestors of birds were estimated to be ~ 163kg around 198 million years ago (Lee et al., 2014), before miniaturisation began, which implies that the appropriate cancer defences must have been in place (analogously to the argument that extant whale life histories would be impossible, based on colorectal cancer alone, if whales did not have better cancer defences than humans; Caulin and Maley, 2011).

Could modern bird life histories be shaped by having inherited pre-existing anti-cancer adaptations from their Mesozoic ancestors, which would mean they may be (i) ‘too robustly built’ for the needs of their current life history, but (ii) simultaneously free to explore, when ecological conditions permit, the life history option of a slow life history? Birds inherit dinosaur genes from very long ago, raising questions about the timescales over which evolutionary lags can operate. In the case of cancer-related adaptations, evolutionary lags occur if gains or losses in cancer defences happen at a slower rate than evolutionary changes in body size and life history (Hochberg and Noble, 2017). We thus examine under what conditions evolutionary lags in cancer-related adaptations are expected to influence extant birds’ longevity and relatively low cancer incidence.

Specifically, we ask: 1) how quickly will defences against cancer adapt when a lineage changes in terms of body size, and under what conditions does this result in evolutionary lags; 2) how do innovations reducing extrinsic mortality (evolution of flight) affect evolutionary lags in cancer-related adaptations? To answer these questions, we model a mutation-selection process over macroevolutionary time scales, spanning several magnitudes of mutation rates. We assume allometric relationships that relate body size to effective population size and extrinsic mortality, which in turn impact the probability of fixation of mutations that alter the expected time that an individual body can resist cancer. Our results show that cancer adaptations can lag behind the body size change in birds under some scenarios. This may result in a longer lifespan and a lower cancer risk, but could also reduce the reproductive success (assuming trade-offs exist between these life history components, i.e. cancer defences are costly; Boddy et al., 2015). On the other hand, innovations that lengthen the average lifespan can select for higher cancer suppression, thereby lowering the effect of these evolutionary lags.

## Results

### Model overview

Here we simulate how cancer defence innovations arise and/or decay in a population via mutations (innovation-decay process, detailed in Methods section *Macroevolutionary simulation*) for 240 million years. We define the relationship between cancer defence and lifespan following the derivation of Kokko and Hochberg (2015). We follow Boddy et al. (2015) in assuming that cancer defences represent a shift towards a slower life history: strong defence yields a (probabilistically) longer lifespan but at a cost of reproductive success per unit time while the organism was alive (implemented as described in Methods section *Deriving the selection coefficient of a mutation*). We assume extrinsic mortality rate scales with body size following the allometric relationships from the literature (McCarthy et al., 2008), and also implement flight as an innovation that can reduce the extrinsic mortality rate in a variant of our model (as detailed in Methods sections *Life history assumptions* and *Modelling body size change and flight*, respectively).

We vary the overall mutation rate across scenarios while keeping the distribution of increases and decreases in cancer defences unchanged. At each time point, we calculate the selection coefficient for each potential mutation by examining how it changes lifetime fitness when the life history of the carrier otherwise follows that established in the current population (body size, extrinsic mortality, presence or absence of flight), but with altered lifespan and reproductive success as these are impacted by the mutation. The fixation probability of cancer-related mutations depends on this selection coefficient, but also on effective population size, which in turn depends on body size based on known ecological relationships. We present an overview of the model in Fig.1 and describe the model details in the Methods section.

**Figure 1:**
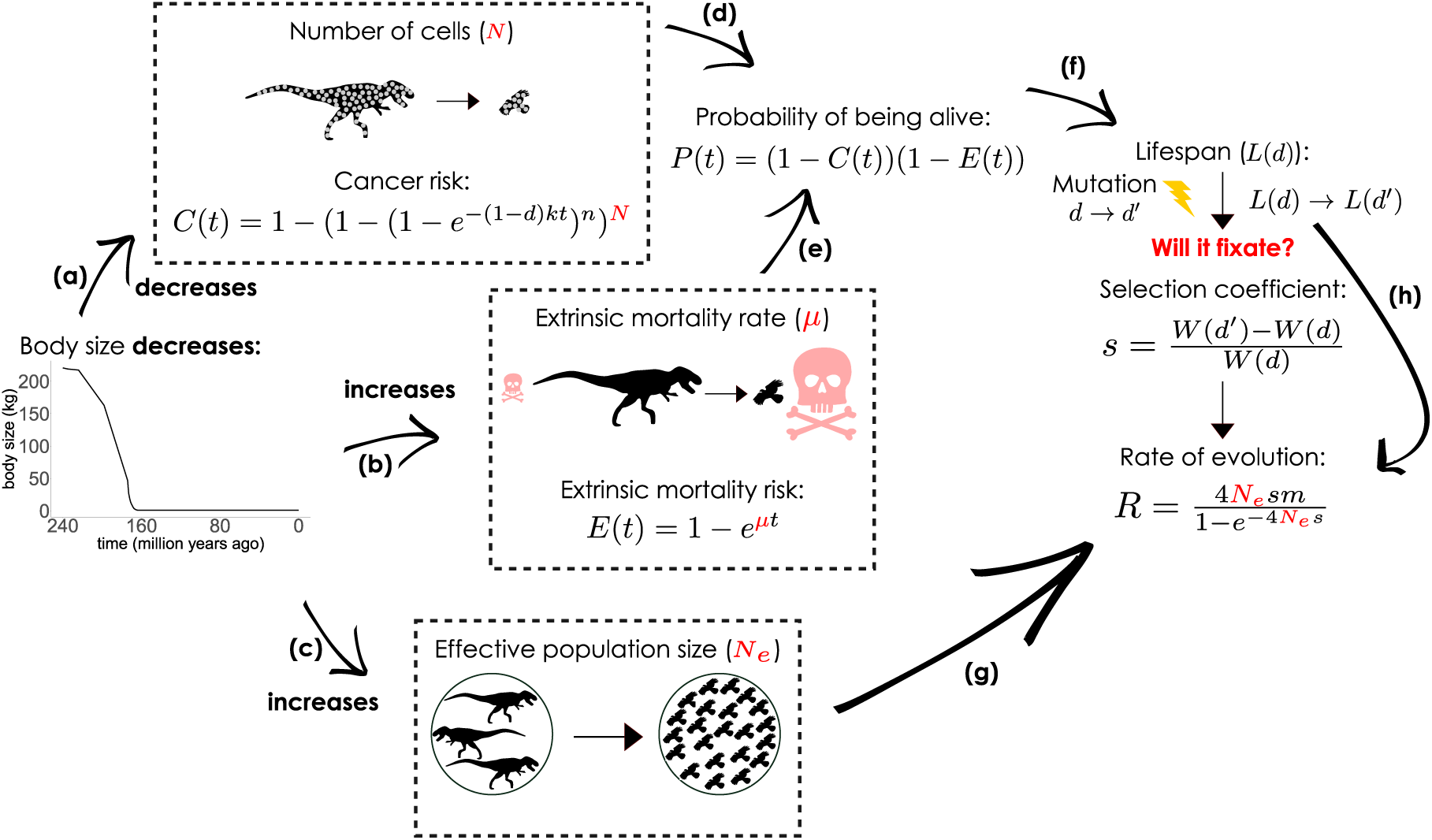
Model overview. Body size, *M* (model notations in Table 1), gradually decreases from 240 Mya to 163 Mya, approximated from Lee et al. (2014), stabilizing thereafter. As the body size shrinks, (a) the number of cells decreases, lowering the age-specific cumulative cancer risk; (b) the ecology changes to increase the extrinsic mortality rate (but with an optional permanent mortality reduction when the innovation of flight happens); (c) population density increases, resulting in a larger effective population size *N*_e_. An individual can reproduce as long as it is alive, the age-specific probability of which is impacted by probability of death (by time *t*) by (d) cancer *C*(*t*) and by (e) extrinsic mortality *E*(*t*), then used (f) to calculate lifespan *L*(*d*). Each mutation is associated with a selection coefficient based on Kokko and Hochberg (2015), (g) which is used together with *N*_e_ to arrive at a rate of this mutation fixing. Mutation rate *m* in our simulations is per genome per generation, therefore, (h) to obtain the rate of at which mutations in cancer defences will arise and fixate (*R*) we use the population’s currently evolved lifespan as a proxy for the generation time.

### Mutation rate and costliness affect the evolution of cancer defences

Mutations in cancer defences did not fixate (grey in Fig.2a) in the population below a certain mutation rate, regardless of body size change or flightedness scenario. We refer to this as the *‘mutational boundary’*. When cancer defences imposed little cost on reproductive success (leftmost part of each subplot), this boundary positioned similarly, just below *m*=10^*−*6^, across different scenarios, except for the constant large flightless lineage where it was *m*=10^*−*5^. Increasing the cost of defences did not affect the mutational boundary for the non-shrinking lineages nor caused any lags in them (Supplementary Information): as these lineages initiated at their optimal defences and did not have any changes in their size or life history, there was no selection to change their level of cancer defences.

**Figure 2:**
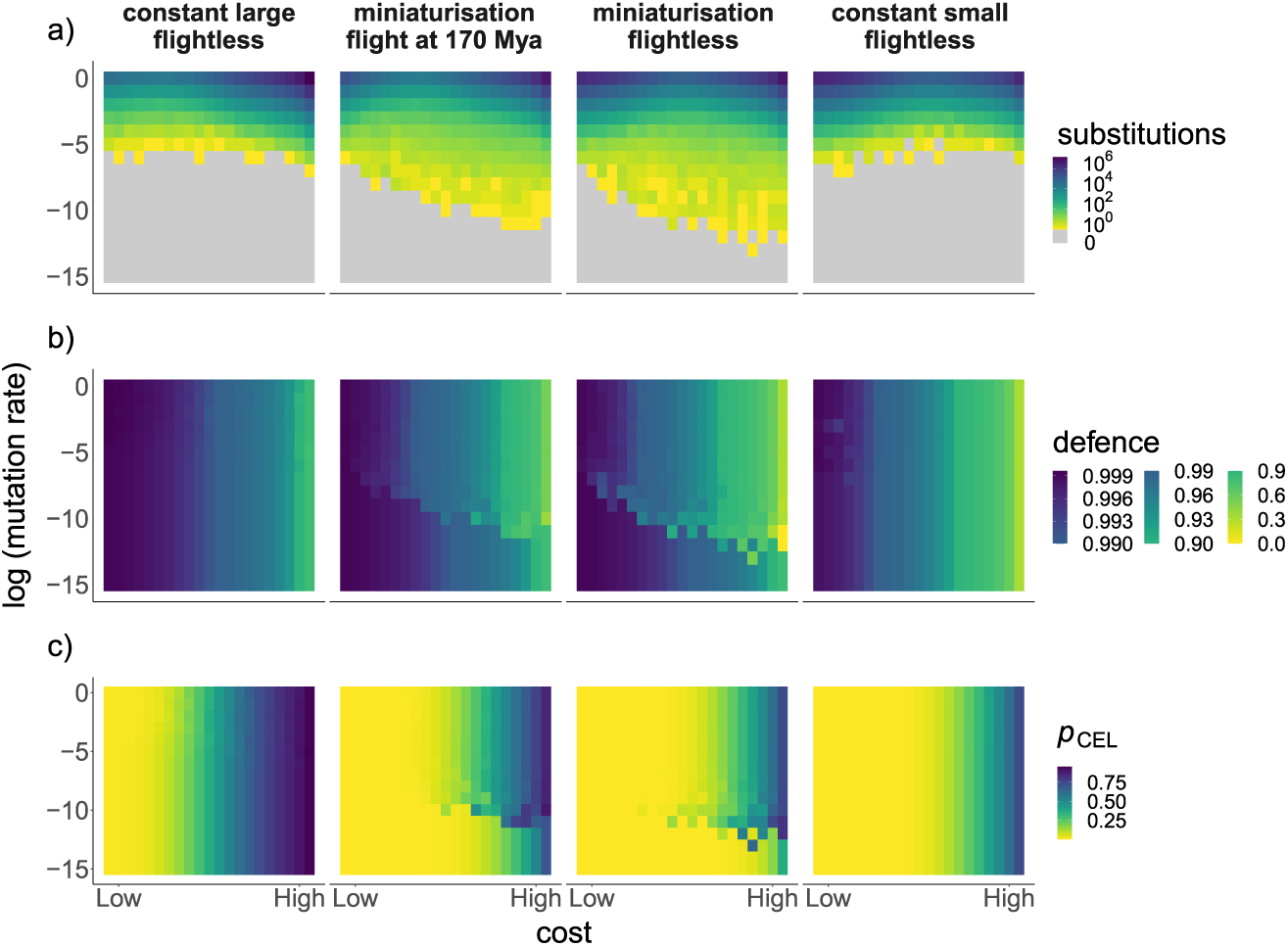
Evolution of cancer defences. Columns refer to different scenarios as indicated. Within each subplot, each pixel combines a specific cost of cancer defence (x axis: log_10_ *c* ranging between −4.1 and −0.1) with a specific mutation rate (y axis: log_10_ *m* ranging between −15 and 0). Rows, top-to-bottom: a) total number of mutations that became fixed by the end of each simulation (grey denotes no fixations), b) level of cancer defences at the end of simulations, and c) probability of cancer ending lifespan (*p*_CEL_). Miniaturised lineages retained ancestral cancer defences and had an overall lower probability of dying from cancer compared to the constant-sized lineages, below the *‘mutational boundary’* (grey area in a). Results are reported for the number of oncogenic steps *n*=3, the rate of oncogenesis *k*=0.0001 for all scenarios, and extrinsic mortality reduction coefficient *r*=1/3 for the flighted case.

Shrinking lineages (the two minituarisation scenarios) experienced a shift between the importance of two different sources of death — extrinsic mortality and cancer — due to body size reduction. Throughout miniaturisation, their extrinsic mortality rate increased (though with modifications for the flighted case, see below) while cancer risk decreased. The resulting selection to reduce cancer defences was negligible if cancer defences were nearly cost-free, but stronger at the costly end of the range that we consider. For the high-cost cases (rightmost part of the subplots), this pushed the mutational boundary to just below *m*=10^−12^ and *m*=10^−10^ for the minituarisation scenarios without and with flight, respectively.

As they shrunk, birds’ optimal level of cancer suppression reduced to become similar to the mammal-like lineage where body size was constantly small and flight did not evolve. Above the mutational boundary, substitutions occurred at a sufficient frequency to make birds’ cancer defences track this shift, such that both flighted and flightless lineages evolved to reduce their cancer defences (Fig.2b). Note, however, that the optimal level of defences when flighted was higher than either of the small flightless lineages (miniaturised and constant-sized).

Below the mutational boundary all lineages kept their ancestral cancer defences. For lineages that maintained their size throughout the macroevolutionary time, this simply meant staying at the optimal level of cancer suppression, with no difference in cancer defences above and below the mutational boundary (constant large flightless and constant small flightless lineages in Fig.2b). In the case of the two bird lineages, where staying below the mutational boundary meant retaining the defences of their large dinosaurian ancestors, cancer suppression remained stronger than in the constant small flightless lineage (indicated with more dark blue in the middle row, or more yellow in the bottom row, of Fig.2), except when cancer defences were assumed to not trade off strongly with other aspects of fitness (low cost), in which case all lineages displayed high cancer suppression, unrelated to other aspects such as flight, body size, or their evolutionary past.

### Low cancer risk and long lifespan at the expense of reproductive success

When costs of cancer defences were low, shrinking lineages displayed little or no evolutionary lags (Fig.3b-d) even when residing below the mutational boundary where ancestral defences were maintained (Fig.2b). Here, the nearly cost-free expression of strong defences meant that being constrained to express high levels of defences (maintained due to phylogenetic inertia) did not incur fitness losses despite the defences being ‘unnecessarily’ strong for small organisms. Here, the probability of cancer ending lifespan (*p*_CEL_, Fig.2c) was similarly low across all body size and flightedness scenarios.

**Figure 3:**
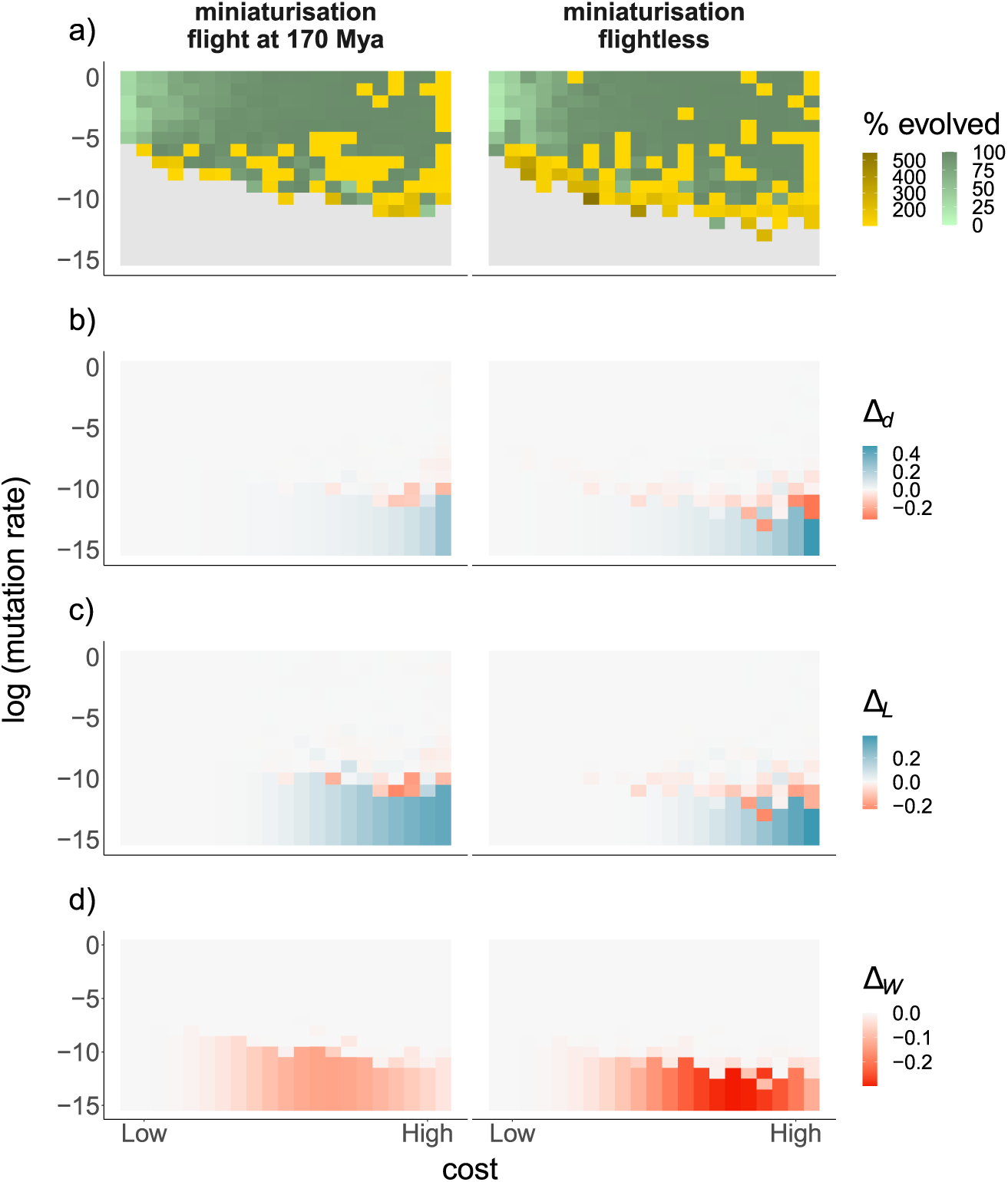
Evolutionary lags at the end of simulations. Columns and panels as described in Fig.2. Rows, top-to-bottom: a) percent of evolutionary change from ancestral to newly optimal defences that is completed by the lineage by the end of the simulation (values *>* 100% indicate ‘overshooting’ the target), lags in b) cancer defences (∆*_d_*), c) lifespan (∆_*L*_), and d) fitness (∆_*W*_). Low enough mutation rates, below the *‘mutational boundary’* (grey) yielded evolutionary lags in cancer defences that permanently lowered the fitness in the miniaturised lineage. Lags that involve higher than optimal values (blue) were prevalent in miniaturised lineages if mutation rates were low and costs were high, but fitness lags (red) were strongest at moderate rather than high costs. The innovation of flight clearly reduced the magnitude of fitness lags. Parameter values are as reported in Fig.2.

The situation changed in the more realistic scenario where the defences were more costly. Now, maintaining ancestral defences became maladaptive even as they continued to prolong lifespan and reduce the probability of dying from cancer. High extrinsic mortality in the miniaturised lineage made individuals typically not live long enough to enjoy the benefits of high cancer suppression, while the costs of suppression were paid throughout the lifetime (reduced reproductive success). In other words, the shrunk lineage was too cancer-robust for its own good. Finally, just above the mutational boundary, we also observe runs in which only very few substitutions got fixed. In some of these cases, cancer defences evolved to be lower than optimal (Fig.3a and Fig.2b), such that fitness evolved to be higher than in the ‘complete lag’ case that occurred when there was no substitutions. Lower than optimal cancer defences can be more adaptive than excessive defences, if the cost-benefit balance shifts to a net benefit because of higher reproductive success of cancer-prone organisms. On the other hand, if a subsequent need to increase the defences emerges, e.g. due to flight, a low mutation rate can hinder innovations to increase the previously-decayed cancer suppression and therefore, can, once more, produce fitness lags (see below).

### Flight innovations reduce the fitness lags

Next, we explored whether flight, by reducing the extrinsic mortality rate in the shrinking lineage, can affect the evolution of cancer suppression. Flight indeed diminished the fitness lags: flying counteracted the otherwise shortening lifespan as body size reduced, and kept the ancestrally high cancer defences useful by delaying onset of cancer to late ages (Fig.2 and Fig.3). The macroevolutionary timing of flight acquisition did not affect the results (Supplementary Information). A smaller *r*, implying a greater reduction in extrinsic mortality rate upon flight acquisition, lowered fitness lags further, but did not fully compensate for the defence lags (Supplementary Information), when we used values in line with those found in the literature (Holmes and Austad, 1994; Healy et al., 2014).

Our results above suggest that innovations that reduce extrinsic mortality allow organisms to live longer and thus diminish the negative fitness effects of ancestrally high cancer suppression. Retaining ancestral defences is one of two ways birds can have evolutionary lags in their cancer defences. The alternative scenario involves birds having already evolved lower cancer defences (suited to a small mammal-like life history, i.e. small-bodied and flightless) before the evolution of flight. This scenario would, for extant birds, predict lower than optimal cancer suppression, but it may be as a whole unlikely because it would combine fast enough evolution (to lower defences) in the preflight era with constraints acting on elevating defences again post-flight. However, given that mutations are more likely to destroy than to elevate existing defences, this remains a possibility, and in our model it is particularly relevant near the mutational boundary, where the evolution of lower than optimal cancer defences was likely (Fig.3a).

## Discussion

Our model outlines the conditions required for a lineage’s evolutionary past to be visible in the bodies of small organisms, in the sense of a legacy of high cancer suppression and consequent long lifespan of birds. Given that defending against rarely occurring conditions may incur suboptimally high costs (a trade-off between probabilistic lifespan shortening and the deterministic costs paid to express the defence regardless of the outcome), extant birds may have adaptively eroded ancestrally high dinosaurian defences, which our model shows to happen under a high mutation rate combined with a sufficiently strong tradeoff. On the other hand, at lower rates that make evolutionary change mutation-limited, cancer-related adaptations may stay in place during the miniaturisation process, and this setting shows higher cancer suppression in birds relative to mammals may reflect birds’ dinosaurian ancestry, potentially explaining the longevity and cancer-resistance observed in birds (Effron et al., 1977; Møller et al., 2017).

Since most small-bodied birds are able to fly, and flight itself may prolong lifespan but could not evolve before avian bodies were sufficiently small to be carried by wings, our model also considers the reversal of selection that may happen as a result of the innovation of flight. From this point onwards, the retention of ancestral cancer defences is again more favourable than it was before, creating nonlinearities in the expected evolutionary lags; here it is relevant that reducing a cancer defence (the pre-flight scenario) may be easier than regaining it post-flight. In such a setting, extant birds would not necessarily be any better defended against cancer than mammals of equivalent body size. At sufficiently high mutation rates, birds could also have gained their own cancer-related innovations which we then predict to be different than those of mammals.

Perhaps counterintuitively at first sight, having evolved or retained better cancer defences do not always translate into a higher fitness: should there be a trade-off between cancer defences and reproductive success, an organism (e.g. a bird) may be ‘excessively cancer-robust’, as reducing defences would allow moving to a faster point at a fast-slow life history continuum, often optimal for small-bodied organisms (Boddy et al., 2015; Brown et al., 2015). Evolutionary lags in our model vanished when reproductive success was not strongly limited by the expression of defences that delay the onset of cancer to later ages.

Our model joins a tradition of others that also rely on a trade-offs between current reproductive success and the ability of an organism to delay the onset of somatic decay (Boddy et al., 2015; Brown et al., 2015), generalizable as a manifestation of antagonistic pleiotropy of ageing (Williams, 1957). Evidence for cancer as a late-life cost of early reproduction is still scarce (e.g. Fernandez and Bowser, 2010; Smith et al., 2012; Werneck et al., 2018), and the strength of this trade-off likely varies between taxa and remains to be better understood (Brown et al., 2015). More generally, trade-offs between longevity and reproductive success are common in nature (see Flatt and Partridge (2018) for a review; Sudyka et al. (2019) for a study showing that reproductive success increases telomere attrition rate in birds). Cancer is merely one way for a body to fail over time, and our model, being suitable to be modified for different failures resulting from insufficient somatic maintenance (Kirkwood, 2015; Webster, 2019), could also shed light on other aspects of senescence.

Our model did not include metabolic rate in any of the intermediate steps between body size and derivations that impact lifespan. Metabolic rate is of potential interest since it has an allometric relationship with body size (Nagy, 2005; Savage et al., 2007) such that larger animals have higher energy expenditure but lower per gram of body mass. It is elevated for flighted birds (compared with flightless ones and mammals; McNab, 2009), with estimates for the timing of increase roughly coinciding with the evolution of flight (Rezende et al., 2020), and a lower cellular metabolic rate has been hypothesized to contribute to cancer risk management of larger animals (Caulin and Maley, 2011; Dang, 2015). These factors would, taken together, imply that minituarization did not only protect birds against cancer: some of the benefits were simultaneously being eroded because of the presumably elevated rates of ageing caused by the evolution of higher metabolic rate. Metabolic rate cannot, however, be straightforwardly equated with cell division rate (a key driver of cancer risk models), and a recent theoretical study suggests that scaling of the metabolic rate alone is insufficient to compensate for the variation in cancer risk that is caused by changing body sizes (Nunney, 2020). As a whole, metabolic rate has been found to be unrelated to longevity in a study of 4100 extant terrestrial vertebrate species (after accounting for body size and phylogeny; Stark et al., 2020), which makes it difficult to incorporate its hypothetical effects into our model.

Evolutionary lags (or examples of maladaptation) occur in nature in numerous systems (Brady et al., 2019). They are often studied in settings where the interests of two entities do not align, preventing (at least) one party from reaching an optimal situation at any time, e.g. the evolutionary arms race between brood parasitic cuckoos and their hosts (Servedio and Lande, 2003), or where environmental changes are so fast that they result in mismatches to the current environment (e.g. high cancer risk in post-industrial humans; Brown et al., 2015; Hochberg and Noble, 2017). We have expanded this view to macroevolutionary time scales, necessitating us to consider very large mutational ranges to reveal when evolution is mutation-limited or not. Mutation rates as a whole are much higher than the lowest range we consider (for birds see Smeds et al., 2016), but only a small subset of all mutations will play a role in traits that modulate cancer risk, and improved defences are yet another subset of those. The exact placement of the mutational boundary will, moreover, depend on our model assumptions (e.g. area used to derive the effective population size from population density).

The pleiotropic assumption that we used — a straightforward trade-off between unidimensional cancer defence level that reduces the rate at which oncogenic steps are completed and an equally unidimensional reproductive success — is also a rather stark simplification of real cancer suppression which requires a complex network of genes (e.g. Trigos et al., 2019). Intriguingly, some of the pleiotropic effects of these genes could effectively constrain the decay of cancer defences, illustrated by the naked mole rat example. These organisms appear to have incorporated burrow-living adaptations into their cancer defences (Tian et al., 2013; Seluanov et al., 2018), allowing one to speculate that should selection for cancer suppression cease to be relevant for them, losing the defence would be selected against if they continue to live underground. It is similarly conceivable that birds’ dinosaurian ancestors had incorporated a trait evolved for a different function into their cancer defence network, and losing that trait in the bird lineage might be challenging if this function still exists and/or primed the evolution of further innovations (Goldberg and Foo, 2020). In line with this idea, several life-history traits, which also affect reproductive success, seem to adapt slowly to changing conditions in passerines (i.e. they show phylogenetic inertia; Pienaar et al., 2013) or birds in general (Cockburn, 2020). The bird-mammal contrast thus offers an exciting opportunity to discover hitherto unknown cancer suppression mechanisms, regardless of whether these have been maintained from dinosaurian ancestors, or represent their own unique innovations like those in the naked mole rats.

## Methods

### Modelling body size change and flight

To disentangle the effect of diminishing body sizes (*M*, see Table 1 for notation) from other evolutionary changes over time, we contrast results obtained with 1) a realistic timeline of body sizes shrinking with time (‘miniaturisation’ scenario) with two hypothetical scenarios that keep body sizes constant either at the 2) ancestral size (‘constant large flightless’, *M* =220.7 kg), 3) final size of the miniaturised lineage (‘constant small flightless’, *M* =0.8 kg) for 240 million years.

**Table 1:** Model notation

| Symbol | Meaning |
| --- | --- |
| $M$ | Body size (in kg) |
| $N$ | Number of cells |
| $\mu$ | Extrinsic mortality rate |
| $N_e$ | Effective population size |
| $t, T$ | time (age of the organism, and year in macroevolutionary time, respectively) |
| $C(t), E(t)$ | Probability of death by time $t$ due to cancer and extrinsic causes, respectively |
| $k$ | Rate of oncogenesis |
| $n$ | Number of oncogenic steps until cancer |
| $d$ | Level of defences against cancer |
| $P(t)$ | Probability of being alive at age $t$ |
| $L(d), W(d)$ | Lifespan and fitness (lifetime reproductive success), respectively, for a given level of cancer defence |
| $c$ | Cost parameter determining the strength of trade-off between cancer defences and fitness |
| $R$ | Rate of evolution; the rate at which mutations that change cancer defences arise and become fixed in the population |
| $m$ | Mutation rate per genome per generation |
| $\tau$ | Time (in generations) until a mutation fixes in simulations |
| $r$ | Extrinsic mortality reduction coefficient |

We do not model selection that led to the miniaturisation of birds explicitly, instead we approximately follow the phenomenologically documented pattern (Lee et al., 2014) from theropod ancestors to a typical extant bird (Fig.1a). We use the dates reported in the text of Lee et al. (2014) with the femur lengths shown in their Fig.1 (converted to body size in kg with the formula in their supplementary material). With PlotDigitizer (http://plotdigitizer.sourceforge.net/), we get additional size and date data from Fig.2b (dataset 1) of Lee et al. (2014). We report all the dates and body sizes in the Supplementary File 1. Hence, in our model, the bird lineage starts at 220.7 kg 240 million years ago (Mya), and reaches the small body size of 0.8 kg at 163 Mya. We assume linear body size shrinkage between any two consecutive dates with body size information. Once miniaturised, we assume the bird lineage not to change in size for 163 million years. We thus ignore extant bird body size variation.

We derive results for our miniaturisation scenario twice: with and without flight acquisition, to examine how a one-off innovation that reduces extrinsic mortality can affect evolutionary lags in life histories. If lower extrinsic mortality prolongs the expected lifespan, then flight might temporarily increase the relative importance of cancer as a cause of death, prompting selection to increase defences. However, this effect may be absent if flight acquisition occurred when cancer defences were lagging behind, being too strong for the current needs in a miniaturizing lineage. This model variant operates like the baseline miniaturisation scenario, except that from the point onwards where flight innovations occur, the extrinsic mortality is multiplied by a coefficient *r* < 1 and thus reduced to a fraction of what the model otherwise assigns to an animal of a specific body size. The associations between size and mortality still operate after flight acquisition. Lee et al. (2014) dates the bird node to ~163 Mya and recent studies suggest powered flight potential in *Archaeopteryx* (Voeten et al., 2018; Pei et al., 2020), as well as other forms of wing-assisted movement in various early ancestors of birds (Pei et al., 2020). Therefore, we chose three time points (180, 170 and 160 Mya) to simulate the initiation of flight.

### Life history assumptions

Various allometric relations exist between body size and physiological and ecological traits (Blueweiss et al., 1978). First, shrinking translates into a lower number of cells *N*, with cell number scaling linearly with body size (Fig.1a). We calculate *N* that corresponds to a given body size *M* using estimates made for humans (Bianconi et al., 2013), *N* = *M*/70 × 3.72 × 10^13^, assuming an average human weight of 70kg. Second, smaller organisms have a higher death rate, mainly due to extrinsic causes such as predation (Fig.1b; Blueweiss et al., 1978). With the relationship between body size and mortality from McCarthy et al. (2008), we calculate the extrinsic mortality as *μ* = *e*^−1.8^*M*^−0.21^. For those time points where flight has already evolved, we multiply this value by a coefficient *r* < 1.

Third, because the fate of mutations with positive or negative selection coefficients depends on effective population size (*N*_e_, details described below in section *Macroevolutionary simulation*), we use known ecological relationships between body size and population density (Damuth, 1981; Juanes, 1986). We assume that miniaturisation allows a species to reach higher densities, while the species range and the relationship between census population size and effective population size are left unchanged (such that we assume no dramatic changes in the mating system and other factors that may impact the relationship between census population size and *N*_e_). Thus, effective population size *N*_e_ is assumed to scale with body size with the same inherent nonlinearity (same value of exponent) as the relationship between body size and density reported by Juanes (1986). This results in a larger *N*_e_ as the body size shrinks (Fig.1c). For our main examples, we use 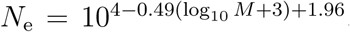, which uses the reported densities for *M* (converted to grams) and derives *N*_e_ by making assumptions about the total range of a species as well as a linear relationship between total (census) population size and *N*_e_. The coefficient 104 corresponds to an area of 10^4^ km^2^ if *N*e equals census population size, or to a larger area if *N*_e_ falls below census population size. Since our choice of species range area is somewhat arbitrary, the relevant check is whether changing the coefficient 10^4^ affects the main results of our study qualitatively; it does not (Supplementary Information).

### Deriving the selection coefficient of a mutation

We track the phenotype ‘level of cancer defence’, *d*, over time, together with its pleiotropic effects on lifespan and reproduction. Mutations change *d* either upwards or downwards, and either type of change may be selected for or against: e.g. a defence-improving mutation may be selected against if maintaining high defences is costly in a situation where cancer is *a priori* rare, such as in very small-bodied organisms. The life history effects of each mutation consist of increased or reduced expected lifespan, as well as the fecundity cost associated with of expressing a specific level of defence. The lifespan effects are modulated by extrinsic mortality: since cancer and extrinsic mortality are two ‘competing’ ways to die, organisms with low extrinsic mortality can reap benefits of high cancer suppression for a longer time than those with high extrinsic mortality, all else being equal (Kokko and Hochberg, 2015).

To compute lifespan for any level *d*, we first follow Boddy et al. (2015) and calculate the probability that an individual with *N* cells, alive at time *t*, has cancer by the time it has lived for *t* units of time, as:

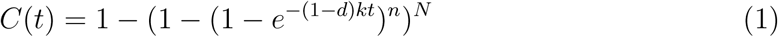

where *d* is the current level of cancer defences, *k* determines the rate at which oncogenic steps occur, and *n* is the number of oncogenic steps that a cell lineage has to complete for the organism to succumb to cancer. The age-dependent increase in cancer risk occurs faster when the number of cells *N* is large, thus defence *d* has stronger effects on lifespan in large than in small organisms. High extrinsic mortality, by virtue of associating with small body size, has a similar effect, with *E(t)*= 1 − *e*^−*μt*^ expressing the probability that an individual is dead by the age *t*. The high mortality *µ* of small-bodied organisms is, optionally, reduced (to an extent) from the time point onwards where flying evolves (as described in *Modelling body size change and flight* above).

An individual is alive at age *t*, i.e. has neither succumbed to extrinsic mortality nor cancer (Fig.1d and Fig.1e), with probability *P* (*t*) = (1 − *E*(*t*))(1 − *C*(*t*)), from which we can derive expected reproductively active lifespan as a function of defences against cancer (*L*(*d*), Fig.1f), as in Boddy et al. (2015). Adaptations to reduce cancer risk can be costly and affect the reproductive output of the organism. We follow Boddy et al. (2015) and model the relationship between fitness (lifetime reproductive success) and cancer defences as *W(d)* = *L(d)*(1 − *d*^1/*c*)^), where *c* is a cost parameter that determines the strength of the trade-off between cancer defences and reproductive success per unit time.

For any current level of defence *d*, we consider an identical mutational distribution of potential changes *δ*: *δ* obeys a generalized extreme value random distribution with parameters *k*=–0.75, *σ*=1, *µ*=–1, which yields values of *δ* between –1 and 1. Positive mutations (*δ* > 0) modify *d* towards the maximum value 1 such that the new value *d′* = *d* + *δ*(1 − *d*), negative ones (*δ* < 0) push *d* towards zero with *d′* = *d*(1 + *δ*), yielding *d′* = 0 if *δ* = −1. These rules combine the plausible assumptions that (i) deleterious mutations (with respect to the level of *d*) are more common than mutations that improve defences, (ii) large increases in cancer defences are rare (for an alternative distribution, see the Supplementary Information), and (iii) the defences as a whole remain between 0 and 1 for the cancer risk equation (1) to make sense. We then calculate the selection coefficient for a given change *δ* in cancer defences as: 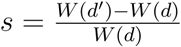 following Kokko and Hochberg (2015).

### Macroevolutionary simulation

We divide the macroevolutionary time into 4000 bins of 60000 years each, assuming negligible body size change within a bin. The number of time bins does not change our results qualitatively (Supplementary Information). The number of generations that occupy one bin is not predefined, as it changes over time due to changes in mean lifespan. Each step in the simulation consists of mutations competing to fix (at rates that depend on the selection coefficient associated with each potential mutation, and the rate at which mutations that diminish or enhance cancer defences occur), determining the next one that fixes, and shifting the macroevolutionary time to the year when the fixing, stochastically, happened.

Each mutation in the mutational distribution associates with a selection coefficient (*s*) as previously derived in section ‘*Deriving the selection coefficient of a mutation*’, with a rate of such mutations occurring determined by the distribution above, and with a consequent rate of fixing *R* based on Lanfear et al. (2014): 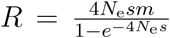, where *m* is the mutation rate per generation, and *N*_e_ is the effective population size. Since *R* depends on the selection coefficient, it is not constant across the different mutations that may arise. We therefore discretize the mutation distribution into 1000 categories by considering mutations that form the 0.05th, 1.5th,…, 99.95th percentile of the mutation distribution, each of them occurring with mutation rate *m*/1000, and having a rate *R*_*i*_ (i = 1, 2,…, 1000) separately derived for each of them according to each mutation’s *s*. While mutations with a high *R*_*i*_ have a better chance to outcompete others (fix before another did), we model the stochastic nature of this competition with a Gillespie algorithm (Gillespie, 1977): the time until a mutation of category *i* fixes is negatively exponentially distributed with mean generation time *τ*_*i*_ = 1/*R*_*i*_; we draw these times for all *i*, and the shortest time found, *τ*_first_, determines the fixation event that actually happens (ahead of any others, which are discarded). Since *τ*_first_ is expressed in the number of generations, we convert this to years by multiplying with the lifespan *L*(*d*) as a proxy for generation time (using the value of *d* before the change in cancer defences) (Fig.1h); Δ*T* = *τ*_first_ × *L*(*d*).

The algorithm then updates macroevolutionary time from its previous value to *T* +∆*T* (*T* being always expressed in years). If the new time is within the same time bin, the post-mutation cancer defence is set to be the new current defence, and the current lifespan is calculated accordingly. If it is not, we instead keep the defence unchanged, update *T* to equal the starting year of the next bin and body size to the pre-assigned size that applies within the new bin. In this case, *L*(*d*), extrinsic mortality rate *µ*, and *N*_e_ are also updated according to the new body size. The entire procedure is repeated as many times as is needed to cover the time from 240Mya to present.

Each body size and flight scenario is initiated with no evolutionary lag at 240Mya (i.e. we assign the level of cancer defences that maximizes the fitness *W* (*d*)). Following Kreyling et al. (2018), we use a gradient design for exploring the parameter space; we simulate each parameter combination without replication, and instead cover a large range with finely adjacent values, as this shows general patterns simultaneously with the degree of inherent variation.

### Evolutionary lags

We quantify the evolutionary lags in fitness, lifespan, and cancer defences, to measure potentially maladapted cancer suppression levels when body size and/or flight ability have evolved differences to ancestral conditions. We first calculate the optimal (fitness-maximizing) level of cancer defences *d∗*, denoting the associated fitness as *W*_max_ = *W* (*d∗*), of the organism for its body size and extrinsic mortality. Similarly, we obtain *L∗* as the lifespan under optimal level of cancer defence. Then, we define the lag in defences as ∆_*d*_ = *d*_final_ *d∗*, where *d*_final_ is the level of cancer defence at the end of the simulations (when *T* =240 million years). Following the same logic, lag in fitness is ∆_*W*_ = log_10_(*W*_final_/*W* (*d∗*)), the ratio between the final fitness *W*_final_ and the maximum fitness *W* (*d∗*), and lag in lifespan is the ratio ∆_*L*_ = log_10_(*L*_final_/*L∗*) (see Table 2 for summary).

**Table 2:**
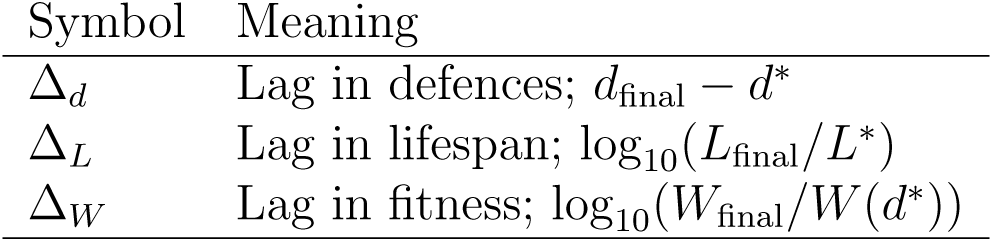
Lag definitions

## Code availability

MATLAB code for the simulations is publicly available in GitHub: https://github.com/yagmurerten/DinoModel.

## Data availability

Raw data generated by the simulations and scripts used for processing and visualization are deposited in figshare, available at https://doi.org/10.6084/m9.figshare.13096664.

## Acknowledgements

This work was supported by the university research priority program (URPP) “Evolution in Action” of the University of Zurich, as well as NIH grant U54 CA217376.

## Author contributions

EYE, MT, and HK conceived the study, EYE and HK developed the model and wrote the paper with input from MT.

