## Supplementary Information for "Bird size with dinosaur-level cancer defences: can evolutionary lags during miniaturisation explain cancer robustness in birds?"

Here we test the robustness of our results for different parameter values and modelling assumptions for: the area size used to estimate the effective population size (Fig.S1), the distribution of mutation steps (Fig.S2), the number of time bins (Fig.S3), and the implementation of flight (Fig.S4). We also expand the Figure 3 in the main text to include lag calculations for the constant-sized lineages (Fig.S5).

### Area size used to estimate the effective population size

In our model, population density scaled with body size and in the main text we used an area of $10^4$ to calculate the effective population size as $N_\mathrm{e} = 10^{4-0.49(\log_{10} M+3)+1.96}$. Here we tested the effect of using different area sizes: $10^3$ (with $N_\mathrm{e} = 10^{3-0.49(\log_{10} M+3)+1.96}$) and $10^5$ ($N_\mathrm{e} = 10^{5-0.49(\log_{10} M+3)+1.96}$).

Using different area sizes changed the range of mutation rates that led to evolutionary lags in fitness (i.e. the positioning of the mutational boundary), but did not otherwise alter our results qualitatively as the evolutionary lags persisted for both area sizes (Fig.S1). Location of the mutational boundary was higher (i.e. at higher mutation rates) for a smaller area ($10^3$) and lower (i.e. at lower mutation rates) for a larger area ($10^5$), compared to the results in the main text (for an area of $10^4$). This is in line with the expectation for an increased strength of selection in large populations (Lanfear et al., 2014), which in our model results in an increased chance of substitutions to reduce cancer defences, and therefore to reduce fitness lags, in the miniaturised lineage.

### Distribution of mutation steps

In our model, we assumed that the distribution of mutational steps reflects the observation that advantageous mutations are rarer than deleterious mutations (Eyre-Walker and Keightley, 2007). For our model this translated into decays in cancer defences being

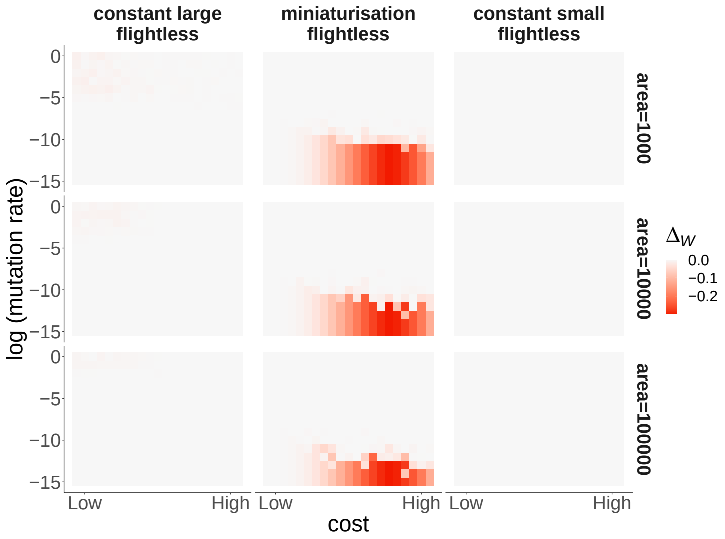


Figure S1: Evolutionary lags in fitness for different area sizes. Each panel shows the cost of cancer defences (x axis: $\log_{10} c$ ranging between -4.1 and -0.1) and mutation rate (y axis: $\log_{10} m$ ranging between -15 and 0). Columns: different scenarios; rows: different area sizes, as indicated in the figure. Middle row shows the area used in the main text. The scale for evolutionary lag in fitness, $\Delta_W$, is shown in the legend. Positioning of the mutational boundary (here inferred from the occurrence of evolutionary lags in fitness) differed, but the results were otherwise qualitatively similar across different area sizes. Results are reported for the number of oncogenic steps $n$=3 and the rate of oncogenesis $k$=0.0001 for all scenarios.

more likely compared to gains. Furthermore, beneficial mutations with large effect were unlikely to happen. For this, we used a generalized extreme value random distribution with parameters ($k$=–0.75, $\sigma$=1, $\mu$=–1), bound between (-1,1). To test the robustness of our results to the distribution of $\delta$, we re-ran our macroevolutionary simulations with mutation steps following a Gaussian distribution with $\delta \sim N(0, 0.3)$, also bound between (-1,1). We observed no qualitative difference in evolutionary lags between the two mutational distributions (Fig.S2).

### Number of time bins

In the main text, we divided macroevolutionary time $T$ into 4000 bins of 60000 years each. In these bins we kept body size and the traits that scale with body size (e.g. effective population size) constant. Changing the number of time bins to a lower (1000) or a higher (16000) value did not affect our results qualitatively (Fig.S3).

### Timing of and extrinsic mortality reduction by flight

In the main text, we reported our results with flight innovation occurring at 170 Mya and reducing extrinsic mortality with the coefficient $r^{-1}$=3. Here we show the results for different timing (180 Mya and 160 Mya) and extrinsic mortality reduction ($r^{-1}$=2 and $r^{-1}$=4) values (Fig.S4). Different flight time scenarios resulted in similar evolutionary lags. In line with our results in the main text, evolutionary lags in fitness shrank with

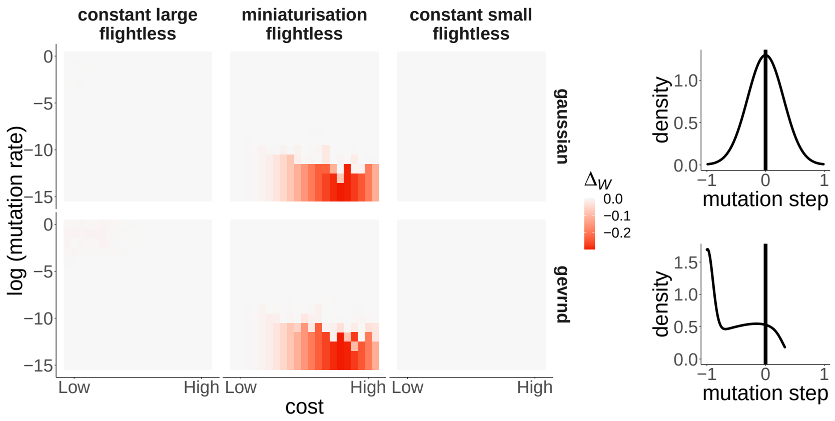


Figure S2: Evolutionary lags in fitness for different mutation step distributions. Panels, columns, and the figure legend as in Fig.S1; rows depict two different mutation distributions: Gaussian (top) and a generalized extreme value (bottom; used in the main text). The evolutionary lags in fitness were similar between the two distributions. Parameters as reported in (Fig.S1), for an area of $10^4$.

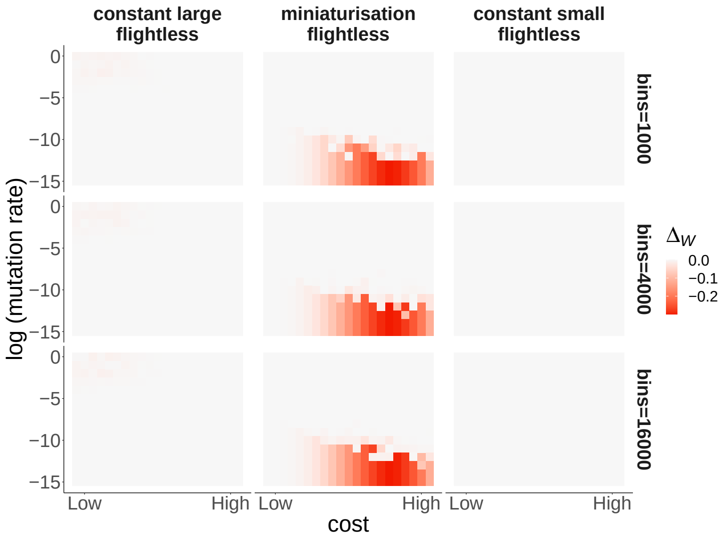


Figure S3: Fitness lags for different numbers of time bins. Panels, columns, and the figure legend as in Fig.S1; rows: the number of time bins. The middle row shows the value used in the main text. Fitness lags and positioning of the mutational boundary were similar across different numbers of time bins. Parameters as reported in Fig.S2.

an increase in $r^{-1}$: reductions in extrinsic mortality rate lengthen the lifespan and allow the 'lagged' lineages to benefit more from their ancestral defences.

### Evolutionary lags for the constant-sized lineages

To complement our results in the main text, here we present evolutionary lags for the constant-sized lineages in addition to the miniaturised ones. As the constant-sized lineages' optimal cancer defences did not change through time (as opposed to the miniaturised lineages that had a shift in their optimal value), they did not experience any lags (Fig.S5)

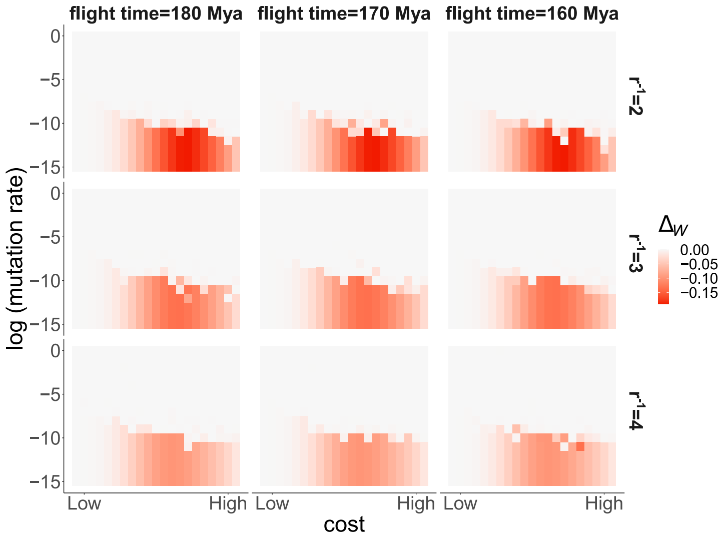


Figure S4: Fitness lags for different flight scenarios. Rows: different extrinsic mortality reduction coefficients ($r^{-1}$; columns: different time points for the flight innovation; panels and figure legend as in Fig.S1. Middle row and middle column (flight time=170 Mya, $r^{-1}$=3) is the parameter combination reported in the main text. The fitness lags became smaller as $r^{-1}$ increased, while the timing of flight acquisition did not affect the results qualitatively. Parameters as reported in Fig.S2.

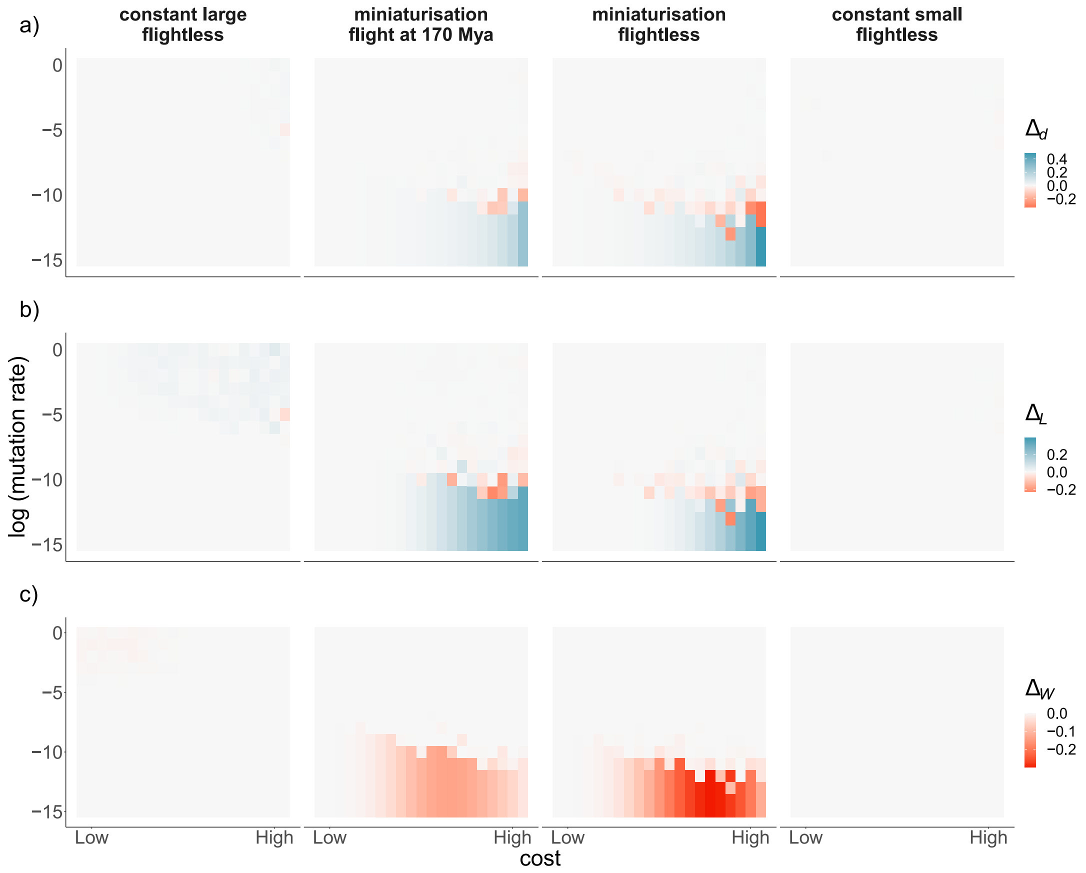


Figure S5: Evolutionary lags for four body size change scenarios. Columns depict different scenarios as indicated, rows show evolutionary lags in a) cancer defences ($\Delta_d$), b) lifespan ($\Delta_L$), and c) fitness ($\Delta_W$). Panels as in Fig.S1. We observe no evolutionary lags in constant-sized lineages. Parameters as reported in Fig.S2.
